# Small molecule inhibitors of hnRNPA2B1-RNA interactions reveal a predictable sorting of RNA subsets into extracellular vesicles

**DOI:** 10.1101/2024.02.05.578546

**Authors:** Jessica Corsi, Daniele Peroni, Romina Belli, Michelangelo Lassandro, Viktoryia Sidarovich, Valentina Adami, Julian Grosskreutz, Fabrizio Fabbiano, Dajana Grossmann, Andreas Hermann, Gianluca Tell, Manuela Basso, Vito Giuseppe D’Agostino

## Abstract

Extracellular vesicles (EVs) are cell-secreted membranous particles contributing to intercellular communication. Coding and non-coding RNAs are widely detected EV cargo, and RNA-binding proteins (RBPs), such as hnRNPA2B1, have been circumstantially implicated in sorting vesicular RNAs. However, the contribution of competitive RBP-RNA interactions responsible for RNA-sorting outcomes still needs to be deciphered, especially for EV-RNA interference and predictability. We conducted a reverse proteomic analysis that prioritized heterogeneous nuclear ribonucleoproteins recognizing purine-rich RNA sequences representing a subset of previously identified EXO motifs. A screening campaign using a full-length human hnRNPA2B1 protein and artificial purine-rich RNA brought to small molecule inhibitors orthogonally validated through biochemical and cell-based approaches. Selected drugs effectively interfered with a post-transcriptional layer impacting secreted EV-RNAs, reducing the vesicular pro-inflammatory miR-221 while counteracting the hnRNPA2B1- or TDP43^Q331K^-dependent paracrine activation of NF-κB in EV-recipient cells. This study demonstrates the possibility of predicting the EV-RNA quality for developing innovative strategies targeting discrete paracrine functions.

**Summary:** Extracellular vesicles (EVs) are cell-released, heterogeneous lipid particles conceived as vehicles for intercellular communication. RNA is a widely detected cargo, and the comprehension of EV-sorting mechanisms represents a step forward in predicting EV quality and associated paracrine effects. While it is known that specific RNA-binding proteins (RBPs) play a role in EV-RNA sorting, the quantitative contribution of competing RBP-RNA interactions and the predictability of RNA-sorting outcomes are poorly understood. Here, we show that a core of hnRNPs compete for the binding to the heterogeneous EV-RNA *in vitro*. Given prioritized interactions with purine-rich RNA motifs, we set up a pharmacological screen platform to find inhibitors of protein-RNA interactions. Our results suggest that selected small molecules can interfere with EV-RNA quality, altering the distribution of specific miRNA cargoes and associating with a discriminant NF-kB activation in EV-recipient cells. This work highlights the role of RBP-RNA interactions in influencing the EV-RNA quality and paracrine functions.

## Introduction

RNA-binding proteins (RBPs) orchestrate different RNA processing activities, from transcription, splicing, maturation, transport, and translation to RNA degradation. Recently, RNA secretion through extracellular vesicles (EVs)^1,2^ has increasingly been recognized as part of the conditional microenvironment in cancer and neurodegenerative diseases^3,4,5,6,7^. EVs are conceived as vehicles of intercellular communication^8,9^. EVs are cell-secreted lipid particles presenting size and molecular diversity due to heterogeneous protein, nucleic acid, and metabolic cargoes^10^. According to their biogenesis, EVs are classified as exosomes, microvesicles, and apoptotic bodies. Exosomes are typically less than 200 nm and generate as intraluminal vesicles within endosomes, involving endosomal sorting complex required for transport through (ESCRT)-dependent or –independent mechanisms^11^. Microvesicles (or ectosomes) range from 100 to 1000 nm in diameter and derive from the outward budding and fission of the plasma membrane through mechanisms likely involving calcium and the interplay of ATP/GTP-dependent proteins^10^. On the other side, apoptotic bodies are larger particles formed during apoptosis^12^.

Coding and non-coding RNAs can be detected in secreted EVs^13,14^. Competing RBPs could significantly contribute to selecting EV transcripts, including, for example, miRNAs^15^, demonstrating that selective packaging can occur at an intracellular level for a desired secretion. We recently highlighted that heterogeneous nuclear ribonucleoproteins (hnRNPs) are frequently detected as vesicular protein cargoes^1^. Among them, TDP43, FUS, and hnRNPA2B1 represent molecular hallmarks of amyotrophic lateral sclerosis (ALS), where these RBPs are known for nuclear loss of function, aggregation tendency, and stress granules assembly^16, 17,18^. In association with the diversity and function of bound RNAs, dosage alterations of several hnRNPs were associated with cancer progression and autoimmune diseases, besides neurodegenerative disorders^19,20,21^. Interestingly, hnRNPA2B1 was recently described as a crucial protein involved in the vesicular sorting of miRNAs through the recognition of specific EXO motifs (5′-AGG, 5′-UAG, 5′-GGAG)^22,23,24^, providing circumstantial evidence that post-transcriptional mechanisms can mediate RNA selection and enrichment into secreted EVs. However, the quantitative contribution of competitive RBP-RNA interactions responsible for RNA-sorting outcomes still needs to be understood, especially for EV-RNA predictability.

To start addressing these biological questions, we designed a reverse proteomic strategy tailored to the secreted EV-RNA and protein binders *in vitro*. We found a significant enrichment of hnRNPs, including hnRNPA2B1, presenting several interaction nodes with RAB proteins already implicated in vesicular trafficking. We distinguished purine-rich RNA sequences as a common substrate recognized by the ranked hnRNPs and constituting a subset of previously identified EXO-motifs. We hypothesized that interfering with hnRNPA2B1-RNA interactions at the intracellular level could represent a tool to modulate the homeostasis of several transcripts and their release through EVs, possibly impacting intercellular communication. Taking advantage of biochemical studies characterizing the binding performance of hnRNPA2B1^25,26^, we performed a high-throughput drug screening to find small molecule inhibitors of the interaction between a human recombinant full-length hnRNPA2B1 and a synthetic purine-rich RNA. We found six hit compounds that efficiently inhibited the interaction *in vitro*, with some passing a validation journey reaching the EV-RNA. By exploiting miR-221 as an elective readout, we observed that Hematein could interfere with the vesicular enrichment of this transcript irrespective of the secreted EV abundance. Preliminary association data indicated that the drug counteracted the hnRNPA2B1- or TDP43^Q331K^-induced NF-κB activation in EV-recipient cells, demonstrating that EV-RNA quality can be modulated by interfering with post-transcriptional control and exploited to develop therapeutic strategies for paracrine functions.

## Results

### Reverse proteomics reveals a competition of heterogeneous nuclear ribonucleoproteins for secreted vesicular RNA

Several RNA-binding proteins, such as hnRNPA2B1, were circumstantially implicated in vesicular RNA (EV-RNA) sorting, given the recognition of specific consensus sequences detectable in vesicular transcripts^22,23,1^. To probe cumulative protein interactions with steady-state EV-RNA, we designed a reverse proteomic strategy^27^ using heterogeneous EV-RNA fragments to pull down intracellular protein binders *in vitro*. By this approach, we assumed that selected RBPs competitively recognize eligible RNAs involving candidates guided by the EV-RNA quality. We optimized the capture probe by hybridizing EV-RNA fragments with a 5’-biotinylated poly(T) oligo, subsequently incubated with Streptavidin-beads to pull down protein-RNA complexes. To normalize the relative protein abundance in cell lysates, we set the same EV-RNA probe in parallel with lysates from cells expressing or not a recombinant hnRNPA2B1 protein and applied mass spectrometry (MS) to identify binding partners resulting from a ratio between the two lysates (**Figure 1A**).

**Figure 1.**
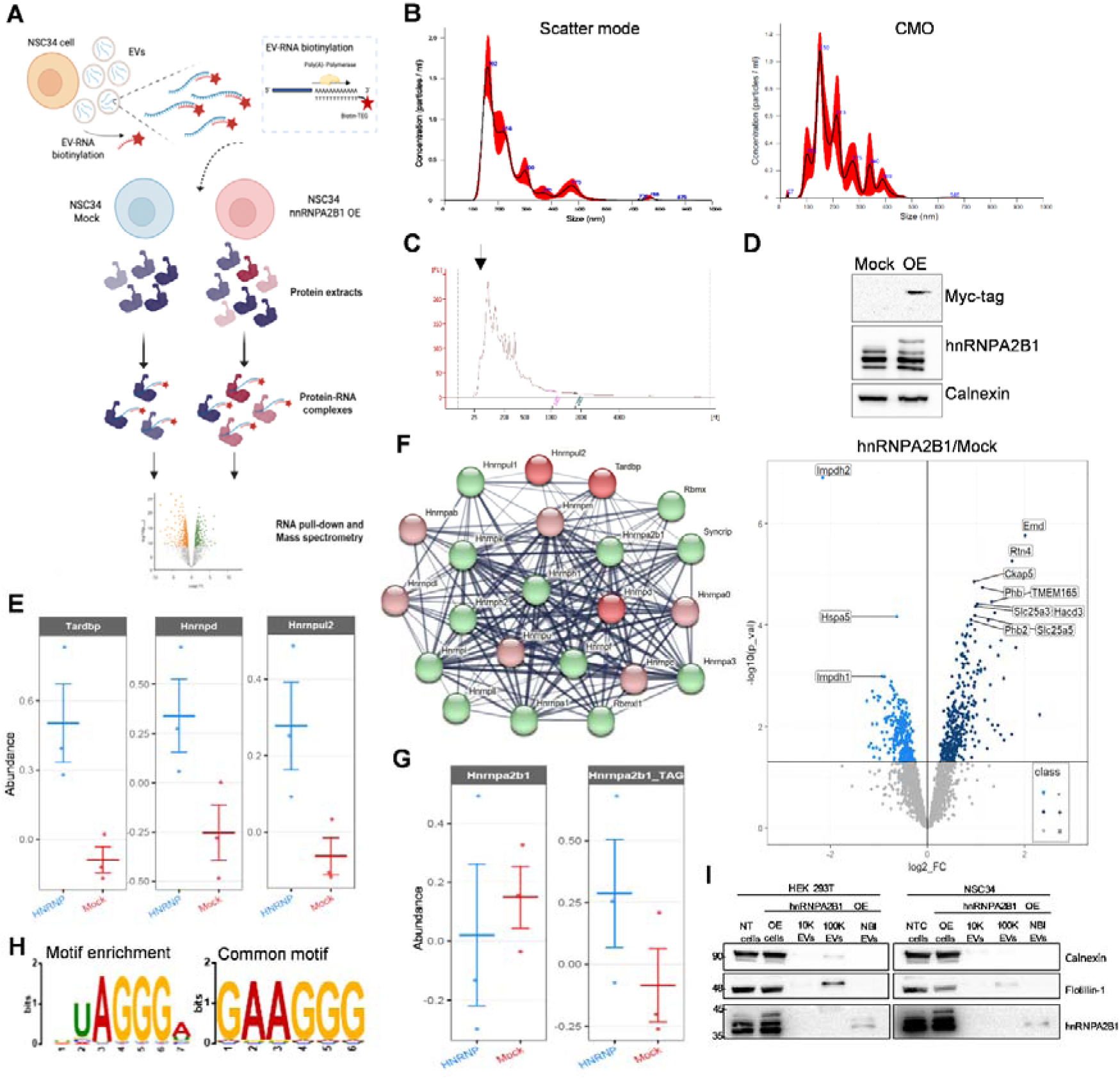
Reverse proteomics using the EV-RNA to prioritize RBP binding partners. (A) Experimental workflow of the reverse proteomic approach. The RNA was extracted from EVs and then enzymatically polyadenylated. A Biotinylated oligo dT was hybridized and then incubated with magnetic Streptavidin beads to constitute the heterogeneous protein-capturing probe. Streptavidin beads with biotinylated oligo dT alone were included as technical negative controls. The probe was then incubated with cell lysates overexpressing or not hnRNPA2B1. Precipitated proteins were analysed by MS/MS and reported as a ratio between the two different cell lysates and from three independent experiments. The image was created on biorender.com. **(B)** Representative nanoparticle tracking analysis profile of NSC34-recovered particles detected in scatter mode (left) and corresponding lipid particles positive to Cell Mask Orange (CMO) detected in fluorescence mode (right). **(C)** Representative Bioanalyzer profile of the EV-RNA subjected to polyadenylation for the reverse proteomics experiments. The black arrow indicates the enrichment of RNA fragments between 100 and 200 nt expected from vesicular RNAs. **(D)** Immunoblotting on NSC34 lysates expressing (OE) or not (Mock) the Myc-His-hnRNPA2B1 protein, recognized by anti-hnRNPA2B1 or anti-Myc primary antibodies. Below, the volcano plot shows the differentially enriched or depleted proteins detected by MS and reported as a ratio between hnRNPA2B1 OE and Mock lysates. Enriched proteins (Class “+”), depleted proteins (Class “-”), and unchanged proteins (Class “=”) are indicated. **(E)** Relative abundance, in three independent experiments, of selected hnRNPs significantly enriched in the hnRNPA2B1 OE condition. **(F)** Network of experimentally validated interactions among the identified hnRNPs obtained from STRING database (https://string-db.org). Red circles indicate hnRNPs significantly enriched in the ratio OE/Mock; Pink circles indicate hnRNPs showing a trend of enrichment; Green circles indicate hnRNPs showing a trend of displacement. Raw data are available in Table S1. **(G)** Relative abundance of recombinant (Myc-His-hnRNPA2B1) and endogenous (hnrnpa2b1) proteins in the ratio OE/Mock. **(H)** Sequence motifs analysed by MEME toolbox (https://meme-suite.org/meme) using experimentally validated RNA sequences present in RPDB database (http://rbpdb.ccbr.utoronto.ca) and recognized by the identified hnRNPs. **(I)** Immunoblotting on cell and EV protein extracts from HEK293T and NSC34 cell lines. “NT”, non-transfected cells; “OE”, hnRNPA2B1 overexpressing cells. EVs were isolated by NBI and eluted particles were subjected to two steps of differential ultracentrifugation (“10K”, corresponding to 10,000 *g*) and “100K”, corresponding to 100,000 *g*) to enrich with larger or smaller size particles, respectively. Flotillin-1 and Calnexin were chosen as positive and negative markers, respectively, of EVs.

We recovered EVs released from NSC34 cells by nickel-based isolation (NBI)^28,29^ and profiled them by nanoparticle tracking analysis (NTA). As previously reported^29^, we observed polydisperse particles ranging from 100 to 800 nm, with small population peaks comprised between 160 and 220 nm (**Figure 1B**, scatter mode). To assess the vesicular fraction of recovered particles, we repeated NTA after staining with Cell Mask Orange, a fluorescent dye for lipid membranes. We detected about 75% of CMO-positive particles, confirming the presence of EVs with the expected size in our preparations (**Figure 1B**, CMO). We isolated the RNA from approximately 10^12^ EVs, and automated electrophoresis showed heterogeneous RNA fragments with a typical peak between 100 and 200 nt^29,30^ (**Figure 1C**). Subsequently, 0.5 µg of EV-RNA were enzymatically polyadenylated to obtain complementarity with a 5’-biotinylated poly(T) oligo. We incubated this heterogeneous RNA source with Streptavidin-beads and performed a pull-down of protein-RNA complexes at equilibrium, without cross-link steps^31^, in control (Mock) and hnRNPA2B1-expressing (OE) cell lysates (**Figure 1D**, Western blotting) for the MS analysis. From the protein ratio, we expected to observe fluctuations reflecting a higher affinity (enriched proteins) or a competition (depleted/displaced proteins) of candidates upon ectopic expression of hnRNPA2B1.

We found 336 significantly enriched proteins and 279 significantly depleted proteins in hnRNPA2B1 *versus* Mock samples (**Figure 1D**, volcano plot, **and Table S1**). The enriched proteins populated the ontologies of RNA-binding, protein transport, and organelle organization with components of membranes and vesicular trafficking (**Figure S1A**). On the other hand, the depleted proteins mainly described the same ontologies but distinguished proteins involved in the immune response. In the protein-enriched dataset, ∼4% of terms (14 out of 336) belonged to the Rab protein family, which significantly emerged in the first molecular function ontology (GDP binding, *p value*= 4.97*10^-12^) together with other Rab proteins showing a trend of enrichment (n=13) or depletion (n=3) (**Table S2**). The second relevant GO (RNA binding, *p value*=2*10^-11^) was populated by RNA-binding proteins (∼18%), with Tardbp, Hnrnpd, and Hnrnpul2 as remarkably enriched (**Figure 1E and 1F**, red circles). This cluster included other 21 hnRNP members showing a trend of accumulation (**Figure 1F**, pink circles) or instead of displacement following hnRNPA2B1 expression (**Figure 1F**, green circles), either linked by direct or functional interactions. The ectopic Myc-His-tagged hnRNPA2B1 protein worked as an internal control since it could be distinguished from the endogenous Hnrnpa2b1 showing a trend of displacement (*P value*=0.12) (**Figure 1G**). These data indicated a selective dynamic on specific protein subsets, possibly including direct and indirect protein-RNA interactions. Consistently, 49 out of 336 up-regulated proteins (∼14.6%) clustered in a dense interaction network with at least nine reported hnRNPA2B1 interactors, including Tardbp and Hnrnpul2 (**Figure S1B**). Focusing on the quality of the EV-RNA responsible for these protein fluctuations, these interactions could indicate proteins converging on the same transcript, with a competitive displacement of proteins recognizing similar RNA consensus motifs^1,32^. To better understand this aspect, we first repeated the RNA pull-down on NSC34 cell lysates upon ectopically expressing a recombinant TDP43, one of the top-ranked hnRNPs (**Figure 1E**). Remarkably, MS results showed TDP43 among the top binders, together with factors that again populated RNA-binding and vesicular trafficking ontologies, but also specific mitochondrion organization components known to be associated with TDP43 function^33,34^(**Figure S1C and Table S3**).

These data confirmed the validity of the reverse proteomics approach and the imposing association dynamics of this protein and hnRNPs on the EV-RNA probe *in vitro*. Subsequently, to get insights into substrate preferences of the identified hnRNPs, we searched for experimentally-validated RNA sequences using the RPDB database^35^. We collected 129 sequences and performed a motif discovery and enrichment analysis using the XSTREME (MEME suite 5.5.1) toolbox. We obtained 20 RNA motifs between 6 and 15 nt in length, with the UAGGGA motif resulting as the most enriched (*p value*=1.78*10^-3^) and the GAAGGG motif as the most common among the hnRNPs (*E value*=3.02*10^-1^) (**Figure 1H and S1D**). Notably, these purine-rich motifs distinguished one of the EXO motifs previously identified in miRNAs enriched in secreted EVs and bound by hnRNPA2B1 in T cells^22^.

In our EV preparations, the hnRNPA2B1 protein could be detected by immunoblotting in small EV fractions secreted by NSC34 or HEK293T cell lines (**Figure 1I**), in contrast to TDP43, not detected in the same conditions (data not shown). Albeit the technical limitations potentially associated with oligo hybridization, EV-transcripts heterogeneity, and affinity of individual proteins, hnRNPs constitutes a dynamic network competing for a plethora of transcripts that are secreted as cargo of extracellular vesicles.

### Human recombinant hnRNPA2B1 binds to purine-rich ssRNAs

To explore purine-rich RNAs as a binding substrate and verify the formation of protein-RNA complexes *in vitro*, we designed a corresponding single-stranded RNA oligo (5’-GGGGAGGUUAGGGAGGAGGGGGGUAGGCGCC, or EXOmotif). We took advantage of previous literature to characterize the RNA-binding activity of hnRNPA2B1^25^ and purified a human full-length, GST-tagged protein expressed in *E. coli* cells (**Figure S2A**). The recombinant protein was used in RNA electrophoretic mobility-shift assays (REMSAs) together with a 5’-labelled infrared dye oligo. The artificial RNA ligand generated detectable protein-RNA complexes and protein oligomerization (see arrows in **Figure 2A**), independently from the GST tag and partially super-shifted by an anti-hnRNPA2B1 antibody. Next, to gain more quantitative and specificity data on this interaction, we set up AlphaScreen assays with a biotin-labelled version of the synthetic RNA, including two shorter RNA oligos (RNA 114: 5’-AAGGACUAGC and RNA 276: 5’-AGGACUGC) already characterized for the affinity with an hnRNPA2B1 isoform^36^, and an AU-rich element (ARE)^37^ equivalent in size as negative control. The numbers 114 and 276 corresponded to the observed Kd values^36^, therefore we expected a better affinity with RNA 114. We determined the best ligands:beads ratio^38^ (hooking point, **Figure S2B**) and then performed saturation binding experiments (**Figure 2B**). The interaction occurred in the nanomolar range and showed the highest affinity for the EXOmotif probe (Kd=3.4±1.6 nM) compared to the other ligand (RNA 114). RNA 276 and ARE oligos showed near-to-background Alpha counts.

**Figure 2.**
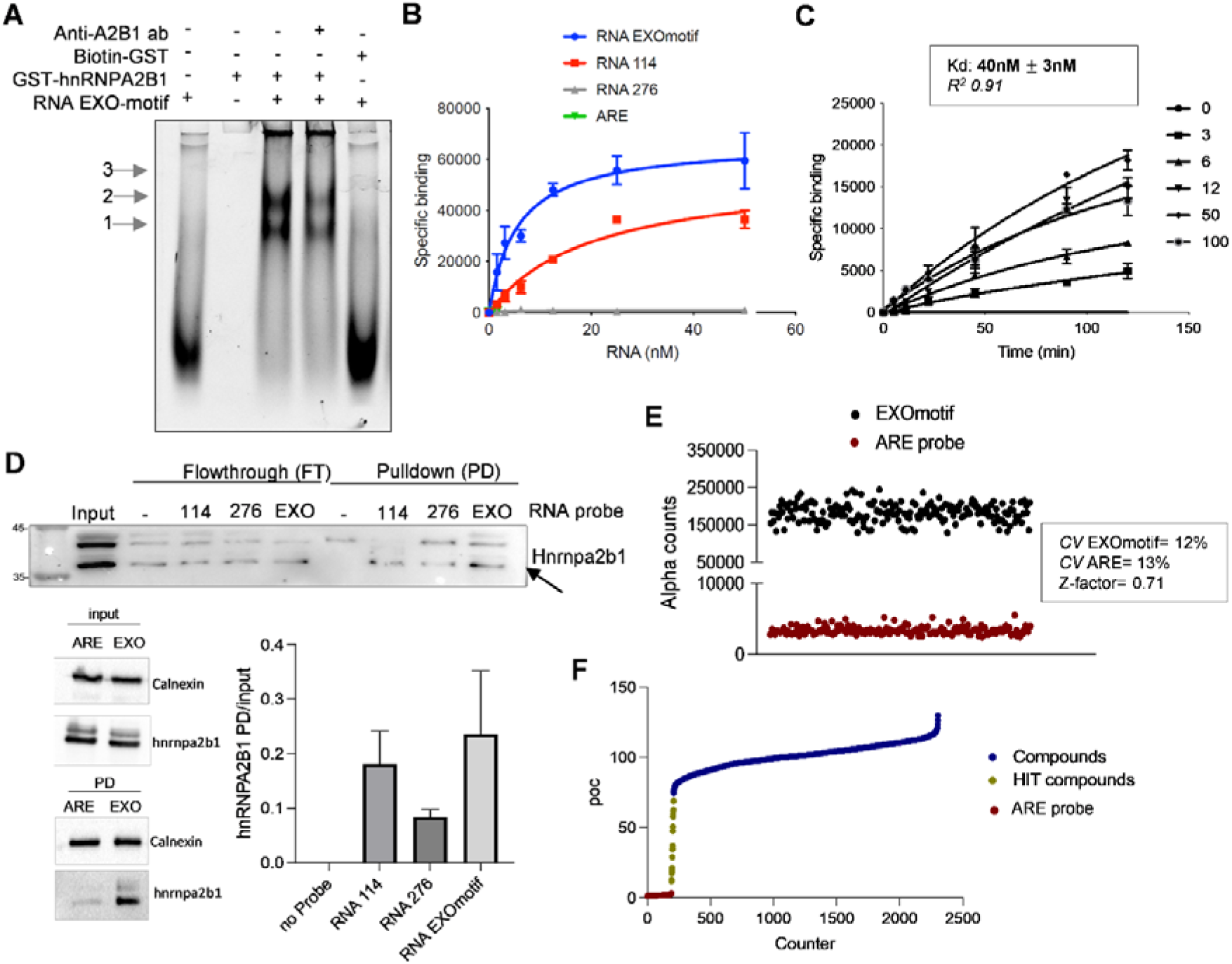
The interaction of recombinant hnRNPA2B1 with purine-rich RNA constitutes a platform for high-throughput drug screening. (A) Representative REMSA performed at equilibrium with 110C:nM of GST-hnRNPA2B1 or Biotin-GST and 25C:nM of RNA probe. Arrows indicate protein-RNA complexes with increasing molecular weights and protein oligomerization phases on the same RNA molecules. **(B)** Representative plot of saturation binding AlphaScreen experiments to measure the GST-hnRNPA2B1 binding to different biotinylated RNA probes, as described in the text. Dissociation constants (Kd) were determined from nonlinear regression 1-site binding model in GraphPad Prism®, version 9.0. Mean and standard deviation values derive from two independent experiments with separate GST-hnRNPA2B1 protein purifications. **(C)** Kinetic experiment carried out with 30 nM of GST-hnRNPA2B1 and increasing concentrations of RNA EXOmotif. Association (*Kon*) and dissociation (*Koff*) rate constants and equilibrium dissociation constants (*koff/kon*) were determined using nonlinear regression association kinetic model of multiple ligand concentration in GraphPad Prism®, version 9.0. Mean and standard deviation derive from two independent experiments with the two protein purifications used in C. **(D)** Representative immunoblotting with anti-hnRNPA2B1 antibody following EXOmotif-, RNA 114-, and RNA 276-based RNA pull down. These experiments were paralleled by an immunoblot showing the higher affinity of the EXOmotif RNA probe compared to the ARE probe. Densitometric quantification of protein fractions were normalized to the input. Mean and SD derive from three independent experiments. **(E)** Distribution of positive (EXOmotif) and negative (ARE) protein-RNA probe interactions calculated in the primary drug screening. Relative coefficient of variation (CV) and Z-factor value are indicated. **(F)** Plot of compounds ranked according to percent of control (POC) normalized (DMSO) values.

Binding kinetic experiments with the EXOmotif probe, in the time frame of 2 hrs, indicated both association (*kon*=523664 M^-1^min^-1^) and dissociation (*koff*=0,02091 min^-1^) rate constants with an observed Kd (*koff/kon*) of 40±3 nM (**Figure 2C**). Since the Kd values between saturation and kinetic experiments significantly differed (3.4 *vs* 40 nM), we ascribed this discrepancy to a protein oligomerization phase impacting the interaction dynamics. Oligomerization of hnRNPA2B1 was already reported^39^ and we observed it by EMSA (**Figure 2A**) and AlphaScreen with a calculated Hill coefficient of 0.3 (**Figure S2B**). Besides this protein behaviour, these results strongly supported the purine-rich sequence as a preferential substrate for the protein. Therefore, we further explored the competitive recognition of this RNA probe by RNA pull-down experiments. We incubated the biotinylated RNA probes with NSC34 native lysates and recovered protein-RNA complexes by Streptavidin magnetic beads. The subsequent immunoblotting showed the precipitation of the endogenous Hnrnpa2b1 protein, displaying multiple bands likely corresponding to post-translationally modified isoforms^40^, only in the presence of RNA probes. In these assays, all the three probes worked with the EXOmotif>RNA 114>RNA 276 affinity ranking, although a modest variability was observed with the purine-rich sequence (**Figure 2D**).

In line with the reverse proteomic findings, these data prioritized purine-rich sequences as relevant substrates for at least hnRNPA2B1 at the intracellular level. Therefore, we used the AlphaScreen platform to identify small molecule inhibitors of the protein-RNA interaction, possibly acquiring a tool for perturbing the secreted EV-RNA.

### Pharmacological screening and orthogonal validation of six drug inhibitors of the interaction with purine-rich RNA

We optimized the AlphaScreen assay to obtain the highest signal-to-background (S/B) ratio and challenge the hnRNPA2B1-EXOmotif interaction with small molecules. We tested a library of almost 2000 compounds plus control DMSO with precautions towards the Alpha assay interference, *i.e.*, using a sub-optimal hooking point and 250 nM of compounds^31^. We calculated a coefficient of variation of 12% for the positive and 13% for the negative RNAs, and a Z-factor of 0.71 indicating a good performance of the primary screening (**Figure 2E**). Compounds were ranked by percent of control (POC) and internal references, such as the biotin interfering with Streptavidin-Donor beads, were excluded from the preliminary short list of inhibitors representing 1% of the whole library (**Figure 2F**). Counter-screening of 21 hits by REMSAs (**Figure 3A**) filtered out six small molecules (Methacycline Hydrochloride or MH, Theaflavin Digallate or TD, Hematein or H, Chrysarobin or C, Phenotrin or P, and Aurin Tricarboxylic Acid or ATA) that were subsequently titrated by Alpha assays in dose-response curves (**Figure 3B**). H, P, and ATA showed almost complete inhibition of the interaction with IC_50_ of about 100 nM, 120 nM, and 250 nM, respectively. The six hits were also tested at 250 nM against the RNA 114 probe and all of them showed a degree of interference, being H and TD the most effective inhibitors (**Figure 3D**). Planarity appeared as a common structural similarity feature among the hit compounds, with an interesting chemical space shared by H and a portion of the TD (red arrow, **Figure 3D**).

**Figure 3.**
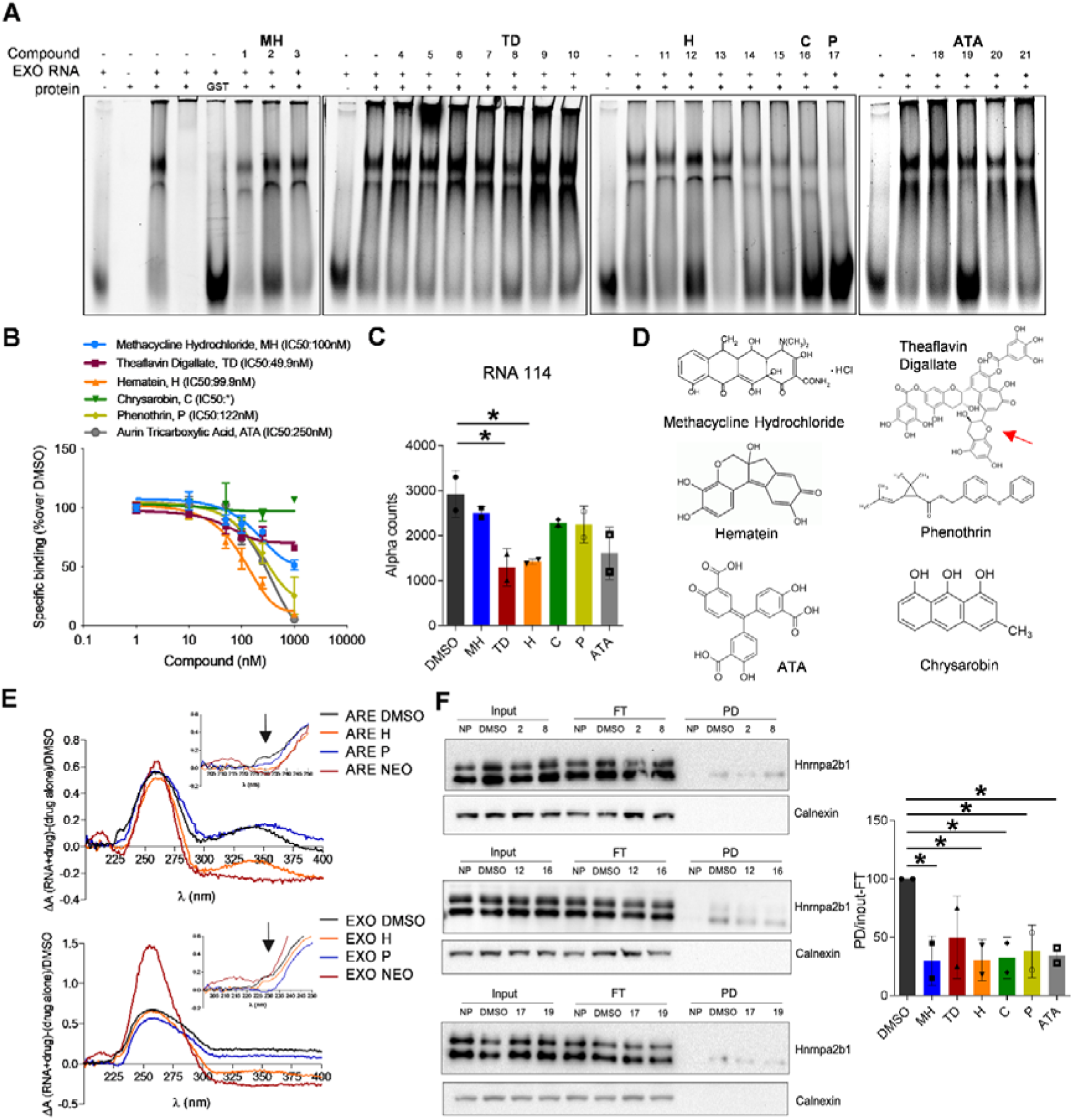
Orthogonal validation of small molecules by biochemical and cell-based assays. (A) Counter screening by RNA lectrophoresis mobility-shift assay (REMSA) showing the activity of AlphaScreen-hit compounds on reducing the hnRNPA2B1-RNA complex formation. **(B)** AlphaScrenn dose-response curves with RNA EXOmotif. The IC_50_ was determined from nonlinear regression using one site fitting model of GraphPad Prism 9. The “*” in the figure indicates “incomplete inhibition”. **(C)** AlphaScreen using 14 nM of RNA 114 probe and 250 nM of compounds. **(D)** Chemical structure of the six AlphaScreen/REMSA hit compounds. Red arrow indicates a chemical space of Theaflavin digallate shared with a Hematein. **(E)** Absorbance spectra relative to Hematein and Phenothrin mixed with RNA EXO motif or ARE probe. Arrows in the inserts magnify the main spectroscopic changes retrieved at 230 nm. **(F)** RNA pull down and immunoblotting of Hnrnpa2b1 using the biotinylated RNA EXOmotif probe upon cell treatment with compounds. Densitometric quantification of the protein is reported as fraction normalized on input-flow-through (FT) and DMSO. * P-value <0.05.

To have an indication of such ligand interference *in vitro*, we acquired absorbance profiles between 200 and 530 nm at room temperature, testing EXO-motif and ARE probes in a 1:5 micromolar ratio with compounds, including neomycin as a positive control^41,42^. We observed spectroscopic changes induced by H and P. In particular, H induced absorbance shifts detectable with both RNA probes and resembling those of neomycin in the range of 300-400 nm, instead slightly impacted by P (**Figure 3E**). At 230 nm, P induced the main spectroscopic changes with the EXO-motif RNA probe, possibly indicating an affinity for secondary structures^41^. These notions were paralleled by a fluorescence indicator displacement (FID) experiment^42^ showing detectable thermal shifts of the EXO-motif RNA of 6.3°C with neomycin, 9.5°C with H, and no detectable shifts with P (**Figure S3**). In the same settings, none of the compounds caused relevant melting shifts of the ARE-containing RNA probe. These data suggested that at least H and P could recognize purine-rich sequences with a certain degree of affinity, possibly depending on primary sequence and/or secondary structures, impacting downstream protein-RNA interactions. This evidence supported further experiments in a cell-based system. Therefore, we treated NSC34 cells for 6 hrs only with 5 µM of compound concentration (far from inducing cell death). The short-term treatment was chosen to minimize potential transcriptional effects induced by the drug, therefore preventing an exacerbated RNA unbalance/adaptation to long-term treatments. After washing with PBS and cell recovering, we applied the same RNA pull-down protocol described in Figure 2D. All the tested compounds significantly prevented the association of hnRNPA2B1 protein to the exogenous purine-rich RNA probe (**Figure 3F**), being MH and H the most effective ones. These data substantiated the possibility of interfering at the post-transcriptional level with purine-rich sequence recognition upon cell treatment.

### Small molecules interfered with secretion of hnRNPA2B1-regulated miRNAs

To measure the extent of perturbation of secreted EV-RNA, we first evaluated the secretome of cells characterized by an altered dosage of hnRNPA2B1. We transfected NSC34 cells to express a Myc-tagged hnRNPA2B1 protein or silence the endogenous protein by siRNA pools (**Figure 4A**). At 48 and 72 hr post-transfection, respectively, the recombinant protein was detected by immunoblotting, as well as an 80% reduction of protein levels in silenced cells. Corresponding media exposed to those cells were used for EV isolation. The relative concentration of particles detected by NTA was slightly perturbed with no statistically significant changes compared with respective Mock or SCR conditions (**Figure 4B**, left).

**Figure 4.**
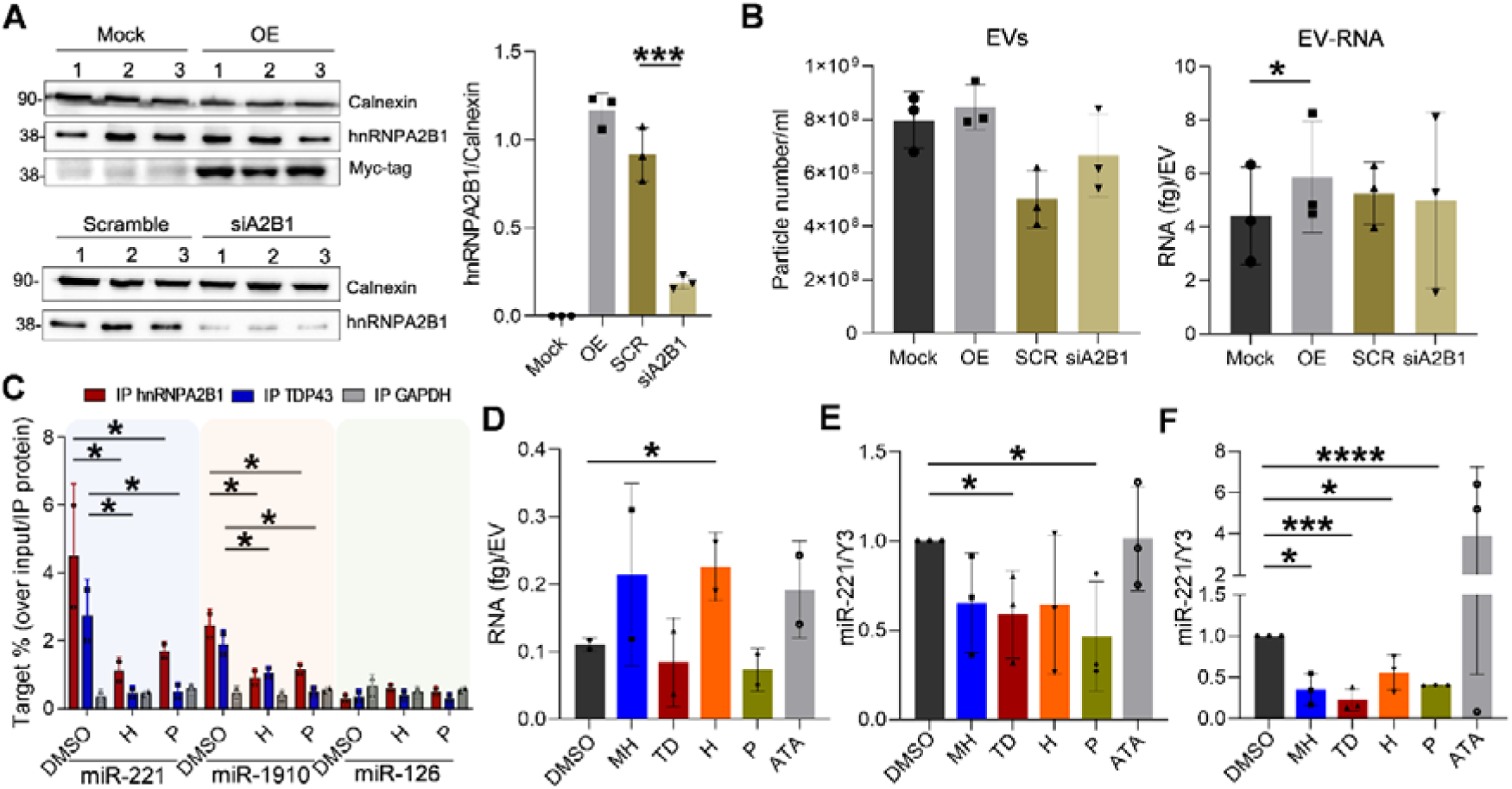
Post-transcriptional activity of small molecules at intracellular and EV-RNA levels. (A) Immunoblotting using protein lysates from NSC34 cells transfected with the plasmid encoding the Myc-tagged hnRNPA2B1 (OE) or without DNA (Mock) and with siRNA pools targeting hnrnpa2b1 (siA2B1) or scramble (SCR). Densitometric quantification of all bands was plotted as a ratio with Calnexin. Mock and OE conditions in the histogram refer to the Myc-tag bands. **(B)** Particle concentration determined by nanoparticle tracking analysis using Nanosight NS300 (left). The EV-RNA concentration determined by Bioanalyzer was normalized on particle numbers upon protein expression or silencing (right). **(C)** RNA immunoprecipitation analysis using cytoplasmic lysates of cell treated with compounds and anti-hnRNPA2B1 –TDP43, and-GAPDH primary antibodies. IgG-conjugated magnetic beads were recovered and divided for protein detection and RNA isolation. Equivalent amount of IP RNA was used for cDNA synthesis and digital droplet PCR with TaqMan probes for miR-221, miR-1910, and miR-126 detection. The transcript copy number was normalized on relative protein levels and DMSO condition. **(D)** Relative abundance of vesicular RNA recovered after cell treatment and normalized on the particle number detected by NTA. **(E)** Relative miR-221-3p copy number detected by ddPCR from NSC34-derived EV-RNA and cDNA synthesis upon compound treatments. Transcript copy number was normalized on Y3 RNA levels and DMSO condition. **(F)** Relative miR-221-3p copy number detected by ddPCR from motor neurons-derived EVs upon compounds treatment. Transcript copy number was normalized on Y3 RNA levels and DMSO condition. * P-value <0.05; *** P-value <0.001; **** P-value <0.0001*.

Further, the EV-RNA profiles of the same samples did not show obvious qualitative changes (**Figure S4A**). However, the EV-RNA abundance appeared highly variable and dramatically fluctuating upon silencing. Nevertheless, we observed a significant trend of accumulation comparing OE with Mock (*p value=0.039*) (**Figure 4B**, right). The observed variability could be consistent with a perturbation of secreted RNA quality, as already demonstrated in hnRNPA2B1-silecing experiments and different cellular models^43,44,22^. Therefore, we decided to perform RNA immunoprecipitation (RIP) analysis on cytoplasmic lysates to verify if compound treatment can interfere with recognition of specific RNA targets at intracellular level. We focused on miR-221 and miR-1910, both harbouring a similar purine-rich motif^22^ in contrast to miR-126. We treated NSC34 cells for 6 hr with DMSO, H, and P (the two most effective compounds). At the end of the treatment, we performed RIP experiments using anti-hnRNPA2B1 and -TDP43 antibodies in parallel with anti-GAPDH as negative control. As shown in **Figure 4C**, both hnRNPs associated with significant enrichments of miR-221 and miR-1910, in contrast to miR-126 and background levels of GAPDH immunocomplexes. Interestingly, the percentage of bound targets was significantly reduced following treatment with H and P, suggesting that compounds can interfere with a selective dynamics of RNA recognition, associated with purine-rich sequences, at intracellular level. Subsequently, we interrogated the EV-RNA. In this case we extended the treatment keeping a 6 hr-schedule with hit compounds. At the end of the treatment, we collected media for EV and EV-RNA isolation. EV-RNA yields appeared highly variable and we observed a significant increase of transcript abundance following treatment with H (**Figures 4D and S4B**).To probe the quality of secreted EV-RNA, we elected miR-221 as a model readout, since it represented an abundant EV-transcript released from different cell types^45,46^, correlated with hnRNPA2B1 dosage^47^, and it was a substrate of several hnRNP members^48^. Interestingly, miR-221 was associated with exosome-mediated tumor phenotypes such as metastasis of cervical squamous carcinoma^49^, malignancy of osteosarcoma cells^50^, and paracrine effects in breast^51,52^ or bladder tumor cells^53^. Notably, miR-221 was one of the inflammatory miRNAs up-regulated in muscles of ALS patients^54^ and a blood-circulating target that positively correlated with progression rate in sporadic ALS patients^55^.

Treatment with compounds MH, TD, H, and P (but not ATA) induced a significant vesicular depletion of miR-221, even after normalization of the miRNA copy number against the endogenous Y3 RNA (**Figures 4E and S4C**).

To validate these notions in other cellular models, we translated the treatment approach to human induced pluripotent stem cell (iPSC)-derived motor neurons^56^. We grew iPSC-derived human small molecule neural progenitor cells (smNPCs) and differentiated them for 24 days to obtain mature motor neurons characterized by Islet1 and ChAt positivity^57^ (**Figure S5**). We treated differentiated cells for 6 hr with compounds and all of them, except ATA, were able to induce a significant down-regulation of secreted EV-miR-221/Y3 copy number compared to DMSO (**Figure 4F**).

In summary, selected bioactive drugs showed a coherent protein-RNA inhibitory activity, encompassing the journey from the biochemical experiments to the cell-secreted EV-RNA, demonstrating that post-transcriptional regulation impact the distribution of specific miRNAs, therefore the quality of EV-RNA sorting.

### Hematein and Phenothrin counteracted the EV-induced paracrine activation of NF-κB

Since miR-221 activates NF-κB in different epithelial cell lines^58,59,60,61^, we studied this paracrine pathway in HEK293T cells, which were transfected with an NF-κB-responsive promoter activating the luciferase reporter. We exposed these cells to EVs isolated from isogenic cells expressing or not the ectopic hnRNPA2B1 protein (**Figure 5A**). The acute treatment with EVs secreted by protein-overexpressing cells caused a significant (*p value*=0.0018) induction of NF-κB responsive promoter expressed as Firefly/Renilla luciferase ratio. Since the S/B improved using transwell co-culture assays (**Figure 5B**), we exploited this system to probe the luciferase activation in EV-recipient cells. By using secretomes from hnRNPA2B1-OE cells treated with small molecules, we found that H could counteract the paracrine, NF-κB-driven activation (**Figure 5C**). Remarkably, additional transwell experiments showed a significant Phenothrin-counteracted paracrine NF-κB activation in wild-type neurons receiving the secretome of TDP43^Q331K^ neurons (**Figure 5D**). In conclusion, this work demonstrates that EV-miRNA secretion is a dynamic circuit composed of connected *cis/trans*-acting factors that small molecules could selectively interfere to challenge EV quality and discrete paracrine functions.

**Figure 5.**
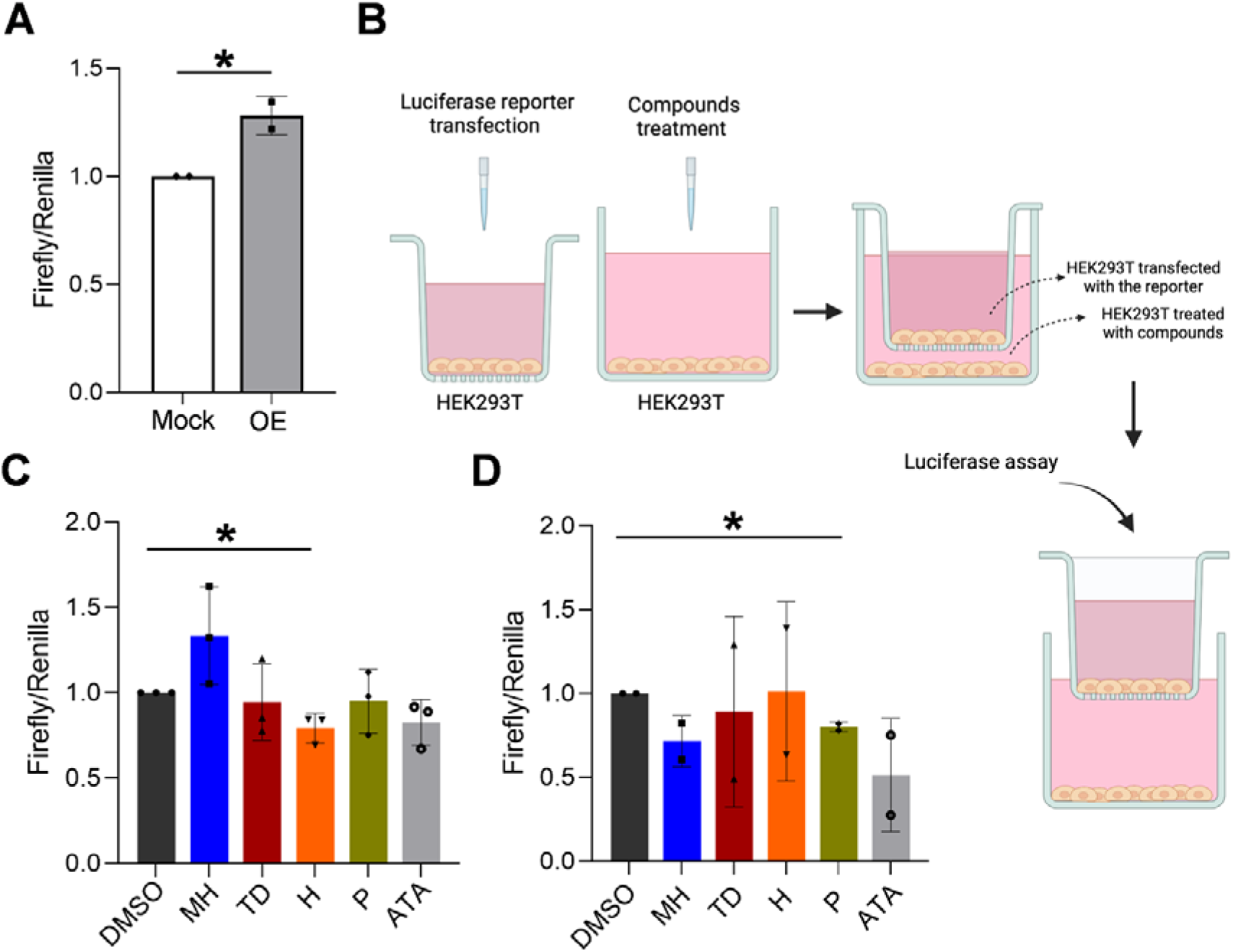
Paracrine effect on NF-κB activation of EVs deriving from RBP-conditioned and treated cells. **(A)** NF-κB activation in HEK293T upon acute treatment with EVs recovered from Mock or recombinant hnRNPA2B1-expressing cells (OE). Firefly luminescence was systematically normalized on Renilla luminescence on the same sample. **(B)** Schematic representation of the transwell setting for NF-κB activation measurement. HEK293T cells were separately plated to treat them with compounds and test them in cells transfected with the luciferase reporter. Cells were washed before exposing them to the co-colture. Image done using biorender.com. **(C)** Transwell experiments to detect NF-κB counteracting effects of compounds in HEK293T cells exposed to the secretome of Mock or hnRNPA2B1-expressing cells (OE). Normalized values were plotted with respect to DMSO. **(D)** Transwell experiments as described in C, but using wild type neurons exposed to the secretome of TDP43^Q331K^ neurons treated with compounds. *P-value <0.05.

## Discussion

In this work, we applied a new biochemical approach to capture cumulative interactions between RBPs and secreted EV-RNA *in vitro*, assuming the “binding” as an exemplified proxy of RNA sorting involvement. Several RBPs have been recently described as part of the EV protein cargo responsible for sorting different RNA species^1^. Since hnRNPA2B1 gained visibility in this process, we considered this protein the first candidate in our perturbation experiments, exploiting the protein enrichment/depletion as a direct surrogate of affinity/competition on the EV-RNA probe. Interestingly, a recombinant hnRNPA2B1 protein representing only 25% of the endogenous protein dosage was enough to detect competition effects, providing evidence of a certain degree of sensitivity. Despite limitations due to a mixture of heterogeneous RNA fragments, which could generate sub-optimal RNA-binding/protein capture, the reverse proteomics approach indicated functional connections between selected hnRNPs and RAB proteins (at least responding to ectopic hnRNPA2B1 expression) over more than 2000 proteins identified by MS. hnRNP members such as hnRNPA2B1 or hnRNPA1 were already described in regulatory networks together with RAB members^62^. Moreover, two ranked hnRNPs (TDP43 and hnRNPD) were already described in direct or functional associations with hnRNPA2B1^63,64^, suggesting possible critical players in concomitant mechanisms of EV-RNA selection and EV biogenesis, yet unclarified.

We prioritized purine-rich sequences by merging the association with hnRNP binders and experimentally validated sequence substrates. Interestingly, the purine-rich RNA was one of the previously identified EXO motifs, strengthening our inference. Further investigation on different cell and EV sources is required to systematically address the relative proportion of secreted RNA sequences. Since TDP43, hnRNPUL2, hnRNPD, and hnRNPA2B1 itself share a common purine-rich RNA recognition, we exploited this notion to set up a platform for challenging RBP-RNA interactions and explore the possibility of interfering with the vesicular distribution of selected transcripts.

We searched for the best strategy to challenge a full-length hnRNPA2B1 protein, widely detected as EV cargo^65^. Unlike the first two common RNA Recognition Motifs, we argued that a full-length protein could behave intracellularly with an oligomerization capacity, virtually affecting the local or general RNA-binding dynamics. Our pilot screening campaign probed small molecules that might present some structural analogies and converge on the inhibition of hnRNPA2B1-EXO-motif RNA interactions. These compounds could biochemically interfere with direct RNA recognition without excluding possible anti-oligomerization effects on hnRNPA2B1. We provide evidence that at least two hit compounds could have RNA-binding selective and/or combined effects in sub-cellular contexts, justifying the observed variable effects on secreted EV-RNA.

The short-term treatment we applied represents a suboptimal schedule not individually tailored to the different cells tested, as we wanted to explore post-transcriptional dynamics, minimizing other secondary events related to cellular stress. Besides their targeting surfaces, the heterogeneous activity of the identified compounds could potentially feature different cell permeability and association rates/modes. The notions we collected here regarding the *in vitro* RNA-binding properties of Hematein and Phenothrin, do not restrict RNA as the only surface interaction of these small molecules. Hematein, for example, has been reported as a protein kinase CK2 inhibitor^66^, while Phenothrin is a synthetic DNA-damaging insecticide^67^. Notwithstanding, these compounds interfere with preferential interactions at the intracellular level and modulate the vesicular RNA sorting, representing new scaffolds in medicinal chemistry to prioritize interactions with RNAs. At this point, other fascinating questions warrant further investigation on other RNA species containing purine-rich motifs, the potential influence of post-transcriptional RNA modifications, and the parallel post-translational modifications on hnRNPs^26,25^.

Since EXO-motif RNA sequences were shown to characterize a plethora of pri- and mature miRNAs, we chose miR-221 as a surrogate readout of EV-RNA fluctuations and a functional target with an established link with pro-inflammation processes. Interestingly, hnRNPA2B1^68^ and miR-221-3p^69^ were individually associated with inflammation. In line with biochemical findings, we could observe a consistent reduction of EV-miR-221 packaging upon short-lasting treatment of different cells. However, the association between the EV biological content and the paracrine NF-κB activation remains elusive, as well as the fate of an EV-mediated horizontal exchange of RNA. We cannot exclude the activation of signalling pathways mediated by EV surfaces or RNA-independent mechanisms, since a set of dedicated experimental strategies is needed to dissect the quantitative contribution of RNA on this pathway. Interestingly, there is already evidence that the reduction of miR-671-5p in EVs from menstrual blood-derived stem cells is responsible for the positive regulation of NF-κB^70^ or that EVs deriving from senescent cells and enriched in miR-30b-5p induce IL-1β and IL-6 interleukins with concomitant activation of NF-κB pathways^71^, or that EVs shuttling miR-660 promote breast cancer progression through a KLHL21-mediated IKKβ/NF-κB p65 axis^72^. These observations are in line with an RNA-induced effect in EV-recipient cells. Consistently, hit compounds interfered with EXO RNA recognition and impacted EV-secreted RNA/quality, counteracting the hnRNPA2B1-induced NF-κB activation in human cells and TDP^Q331K^-induced NF-κB activation in primary mouse neurons, irrespective of the number of particles released. This activation is considered pathogenic in ALS^73,74^, and we provide evidence that secreted EVs upon altered physiology of hnRNPA2B1 or TDP43 mediate this paracrine effect. Besides a conceptual drug-repositioning, we demonstrate that a substantial post-transcriptional control impacts selected EV-RNA secretion, and interfering with the underlying protein-RNA interactions can be instrumental in modulating the EV-associated paracrine physiology (**Figure 6**).

**Figure 6.**
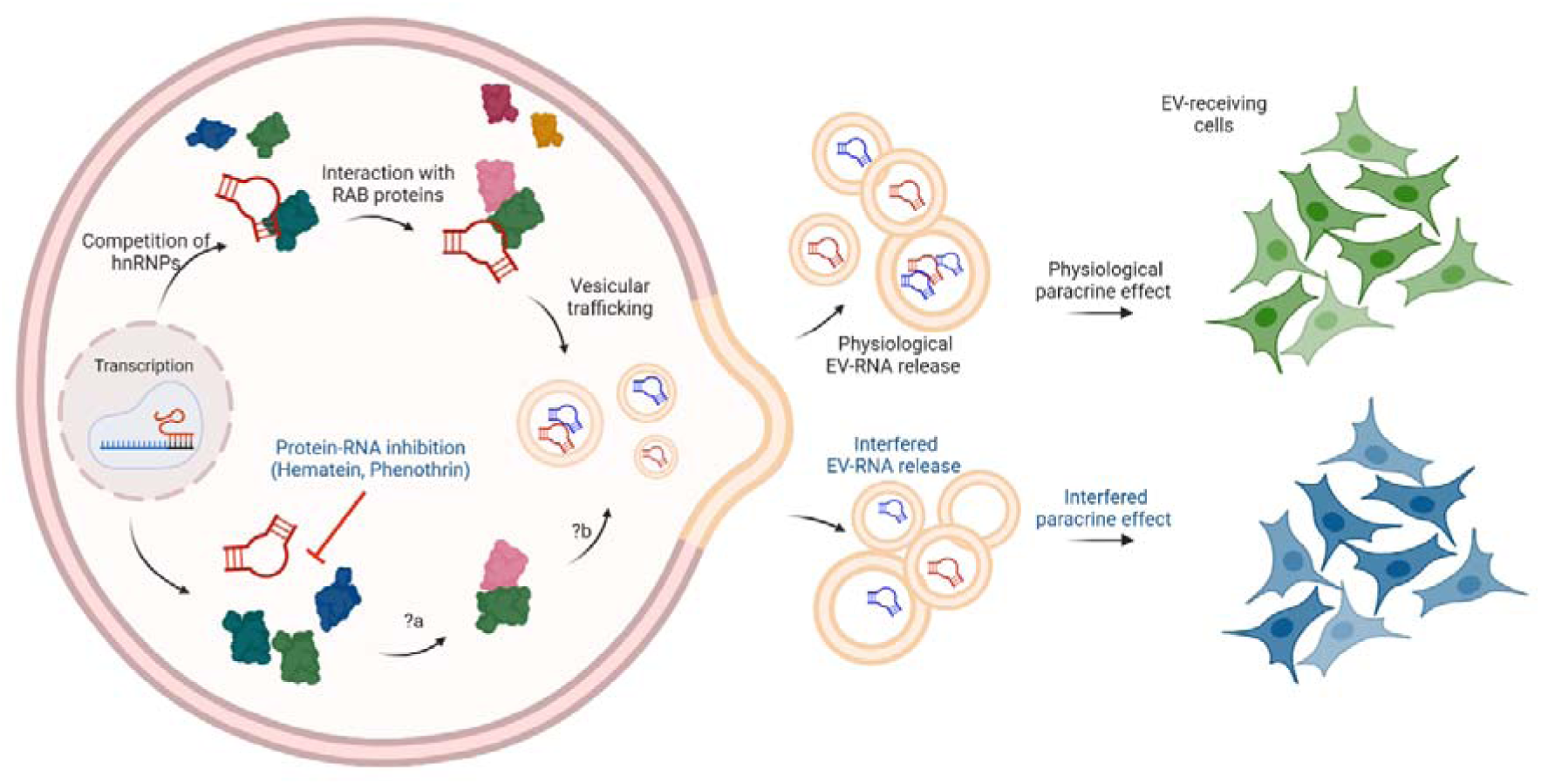
Post-transcriptional control of extracellular vesicle RNA. Schematic representation of cytoplasmic events conveying RNA subsets into secreted EVs. RBPs, and in particular hnRNPs, could compete for and select RNAs. Interactions with other *trans*-acting factors, virtually RAB proteins participating in vesicular trafficking dynamics, could contribute to finalizing the RNA sorting process. Small molecule inhibitors of protein-RNA interactions, such as Hematein or Phenothrin, could alter intracellular RNA homeostasis and, ultimately, the distribution of the secreted counterpart, impacting paracrine functions. Direct consequences following the RNA recognition interference (?a) and vesicular packaging (?b) are still unclear. Image created with Biorender.com.

We open a new scenario where the simultaneous manipulation of RBPs and RNA harboring specific consensus sequences could be explored to identify new drugs with potential clinical applications targeting paracrine functions.

## Acknowledgements

We warmly thank Prof. Sohail Tavazoie (The Rockefeller University) and Prof. Claudio Alarcόn’s group (Yale School of Medicine) for providing us the plasmid to express a GST-tagged hnRNPA2B1 protein.

We thank Dr. Toma Tebaldi for support on statistical analysis of proteomics data.

## Author contributions

VGD and MB conceived the study; VGD designed and supervised experiments; JC performed experiments and statistical analyses; DP and RB performed LC-MS experiments and related statistical analyses; ML participated for a time in pilot protein expression and AlphaScreen set up; JC, VS, and VA co-conducted drug screening and primary analysis; JG and AH provided infrastructure and supervised experiments with motor neuron cells; JC and FF performed transwell analysis; JC and DG performed experiments with iPS cells; GT supervised hnRNPA2B1 interactome; VGD wrote the manuscript and all the authors contributed and revised it.

## Methods

### Cell cultures and reagents

Mouse motor neuron-like hybrid cell line (NSC34) and human embryo kidney HEK293T cells were maintained using Dulbecco’s modified Eagle’s medium (DMEM), supplemented with 10% Fetal Bovine Serum, 1% L-glutamine, and 1% penicillin/streptomycin (Life Technologies, Carlsbad, CA, USA) in standard conditions at 37°C with 5% CO2.

Induced pluripotent stem cells (iPSCs)-derived small molecule neuroprogenitor cells (smNPCs), produced as described in ^57,75^, were differentiated to reach 2-weeks matured motor neurons. The iPSC generation and respective characterizations was previously reported^57^ and approved by the local ethics committee (EK45022009). Briefly, smNPCs were seeded on Matrigel (Corning 354234) coated 12-well plates and maintained in N2/SM1 base medium (made of 48.75% DMEM/F12, 48.75% Neurobasal, 1% Pen/strep/glut, 1% SM1 and 0.5% N2) supplemented with 3 µM CHIR99021, 150 µM Ascorbic acid, and 0.5 µM Purmorphamine (PMA). For the differentiation, smNPCs were plated on Matrigel coated 6-well-plates and fed with N2/SM1 base medium supplemented with 1 µM PMA, 1 ng/ml BDNF, 0.2 mM Ascorbic acid, 1 µM Retinoic acid and 1 ng/ml GDNF for 10 days. Then the cells were plated on 15% Poly-L-Ornithine (Merck/Sigma A-004-C) and 1:100 Laminin-(Biotechne 3400-010-02) coated plates and maintained in N2/SM1 base medium supplemented with 5 ng/ml Activin A (For the first day only), 0.1 mM dBcAMP, 2 ng/ml BDNF, 0.2 mM Ascorbic acid, 1 ng/ml TGFβ-3, and 2 ng/ml GDNF for two weeks. Then, smNPC-derived motor neurons were exposed to 5 µM of compounds for 6 hr.

Primary cortical neurons were cultured from embryonic E15.5 C57BL/6J mice as previously described^76^. TDP-43^Q331K^ mice were conducted as already described^77^ according to institutional guidelines, national (D.L. no. 116, G.U. suppl. 40, February 18, 1992, n. 8, G.U., 14 July 1994), and international laws and policies (EEC Council Directive 86/609, OJ L 358, December 12,1987; National Institutes of Health Guide for the Care and Use of Laboratory Animals, US National Research Council, 1996).

### hnRNPA2B1 over-expression and silencing

NSC34 and HEK293T cells were transfected with a plasmid coding for the hnRNPA2B1 (NM_002137) Human Tagged ORF Clone (Origene, RC219318) using Lipofectamine™ 3000 Transfection Reagent (Invitrogen L3000001). After 48 hr of transfection, cells were used for the subsequent analysis. Protein silencing was obtained using siRNAs targeting *Hnrnpa2b1* or *hnRNPA2B1* genes (ON-TARGETplus mouse Hnrnpa2b1 siRNA-L-040194-01-0005; ON-TARGETplus human HNRNPA2B1 siRNA-L-011690-01-0005). Scramble (SCR) siRNA (ON-TARGETplus Non-targeting siRNA #3, 5 nmol Catalog ID: D-001810-03-05) was used as a control. INTERFERin® (Polyplus-101000036) was used as a transfection reagent in this case.

### Recombinant protein expression and purification

pGEX-6P-1-HNRNPA2B1 plasmid was used to transform BL21(DE3) competent cells. A pre-inoculum of transformed BL21(DE3) bacteria cells was grown in LB broth (Sigma-Aldrich) supplemented with Ampicillin (50 μg/μl) ON at 37°C at 220 rpm. The day after, bacteria were diluted in 1L LB supplemented with 50 μg/μl Ampicillin and cultured at 37°C at 220 rpm. When optical density at 600 nm reached 0.7, the protein expression was induced with 0.2 mM IPTG for 13 hr at 25°C. The culture was harvested by centrifuging at 6371 × g for 15 min at 4°C and the pellet resuspended in 30 ml of lysis buffer (50 mM Tris HCl, pH 7.5, 2 mM EDTA, 200 mM NaCl, 1:1000 Lysozyme (stock 25 mg/ml), 1 mM DTT, bacteria protease inhibitor cocktail) and incubated on ice for 20 min. Upon adding 0,1% tween, the lysate was incubated for an additional 20 min on ice followed by sonication (7-8 cycles – 20 seconds ON, 30 seconds OFF). Sonicated samples were centrifuged at 13’000 *g* for 45 min at 4°C. Then the supernatant was filtered (0.45 μm) and incubated with buffer equilibrated Pierce™ Glutathione Agarose beads (Pierce, 16101) for 2 hr at 4°C in rotation. The lysate was loaded in a column and, after the collection of the flow-through (FT), beads were washed with 25 ml of high-salt wash buffer (50 mM Tris HCl, pH 7.5, 500 mM NaCl). GST-hnRNPA2B1 was eluted in 15 ml elution buffer (100 mM NaCl, 50 mM Tris HCl, pH 7.5, 0.1 mM DTT, 300 mM Glutathione) and concentrated using Amicon Ultra-15 at 4000 g (Merck Millipore).

### AlphaScreen and pharmacological screening

AlphaScreen assay was used to study the interaction between hnRNA2B1 and different biotinylated single-stranded RNA probes: EXO RNA (5’-BTeg-GGGGAGGUUAGGGAGGAGGGGGGUAGGCGCC), RNA 114 (5’-BTeg-AAGGACUAGC), and RNA 276 (5’-B-Teg-AGGACUGC). The assays were performed in OptiPlate-384-well plates (PerkinElmer-6007299) in 20 μl final volume. The ligands were diluted in AlphaBuffer (25 mM HEPES pH 7.4, 100 mM NaCl, 0.01% BSA) and tested using AlphaScreen GST Detection Kit (PerkinElmer, 6760603C). For the assay optimization different concentrations of the RNA probes (0-100 nM) were incubated with different concentrations of GST-hnRNPA2B1 (0-60 nM) in the presence of streptavidin Donor beads and anti-GST-Acceptor beads (PerkinElmer) (20 µg/ml final concentration). For the reactions, 4 μl of the RNA probes were firstly added to the plate, then 16 μl of a mix containing all the other components was added and the plate incubated at room temperature for 1 hr in the dark.

The high-throughput drug screening was performed in a total volume of 20 μl using 17 nM of RNA EXO motif (chosen below the hooking point), 30 nM GST-hnRNPA2B1, and 250 nM of compounds belonging to MS Spectrum Collection library (MicroSource). Compounds were dispensed into white 384-well Optiplates (PerkinElmer) using Echo 650 acoustic dispenser (Beckman Coulter) followed by the addition of protein in Alpha Buffer using the EL460 dispenser (BioTek). After 15 min of incubation, RNA probes were added by Echo 650 acoustic dispenser, while Donor and Acceptor beads from the GST detection kit (Perkin Elmer) were dispensed by EL460 instrument. Following 60 min of incubation, Alpha signal was detected using Enspire plate reader instrument (PerkinElmer). The raw Alpha counts were normalized to DMSO-treated controls within every plate.

### RNA Electromobility Shift Assay (REMSA)

hnRNPA2B1 (110 nM) was incubated with compounds (1 μM) in Remsa Buffer (20 mM HEPES pH 7.5, 50 mM KCl, 0.5 µg BSA, 0.25% Glycerol) at room temperature for 10 min. Then, 30 nM RNA EXO were added in a final volume of 20 μl, and the reaction mix incubated at room temperature for 50 min. The mix was then loaded in a 4% native polyacrylamide gel containing 2% glycerol and run in 0.5X TBE buffer at 60V for the first 15 min and at 80V for additional 60 min. The RNA probe signal was detected using Typhoon Instrument (Amersham™ Typhoon™ 5 - 29187191) using filters for infrared emission detection.

### EV isolation and characterization

EVs were isolated from HEK293T, NSC34 and smNPC-derived motor neurons using Nickel-based Isolation (NBI) or Ultracentrifugation (UC). NBI protocol was performed as already described^28^. Briefly, after a first centrifugation step at 2,800 *g* for 10 min, nickel beads were incubated for 30 min with serum-free media collected from cells in a ratio of 25 μl of beads per ml of medium. After a 2 min centrifugation at 600 *g*, particles were eluted from the beads by adding 1X elution buffer made of Solution A (16 mM EDTA, UltraPure pH 8.0, ThermoFisher) and Solution B (10 mM NaCl, 225 μM citric acid, Sigma-Aldrich) diluted 5 times in PBS. After 15 min incubation at 28°C in a thermoshaker, the EVs were retrieved by 1-minute centrifugation at 1,800 *g* and subjected to the following characterization. Differential ultracentrifugation was performed by a SW 32 Ti swinging bucket rotor in an Optima XPN-100 ultracentrifuge (Beckman Coulter, Brea, CA, USA). The cell-conditioned serum-free media, after the first centrifugation step at 2,800 *g* for 10 min, was put in open-top Ultra-Clear centrifuge tubes (Beckman Coulter-344058) and subjected to 100,000 *g* ultracentrifugation for 70 min at 4 °C under vacuum. Then the supernatant was removed, and the pellet was resuspended in 0.22 μm filtered PBS. The retrieved particles were then used for the subsequent experiments.

NTA was applied for particle characterization using NanoSight NS300 instrument (Malvern Panalytical Ltd., Malvern, UK) equipped with a 488 nm blue laser and a sCMOS camera. EVs samples were diluted in PBS in order to have at least 20 particles/frame and three individual consecutive 60 seconds videos, using 14 as camera level were performed. Information about particles concentration, mean and mode diameter were retrieved using built-in NanoSight Software NTA3.3.301 (Malvern).

### RNA immunoprecipitation (RIP)

Approximately 2*10^6 cells were used for each RIP experiment, performed as already described^31^, without cross-linking steps and using 0.5C:μg/ml of antibodies. Beads-precipitated samples were divided for immunoblotting or TRIzol-based RNA isolation. Densitometric analysis obtained using Image J software (v.1.54). miRNA TaqMan probes were bought from ThermoFisher with the following codes: miR-221 (477981), miR-1910 (479581), and miR-126 (4427975).

### RNA extraction and cDNA synthesis

RNA from cells was extracted using TRI Reagent® (T9424, Sigma) following the manufacturer’s instructions. ThermoFisher Nanodrop 2000 Spectrophotometer was used to assess the purity and quantity of extracted RNA. RNA from EVs was extracted using Single Cell RNA Isolation Kit (51800, Norgen) following the manufacturer’s protocol. The elution was in 10-20 µL of RNase-free water and the RNA profiling and quantification was assessed by Agilent 2100 Bioanalyzer and RNA 6000 Pico Kit (5067-1513). Half nanogram of extracted RNA from cells and EVs was used as input for cDNA synthesis. cDNA was produced using TaqMan™ Advanced miRNA cDNA Synthesis Kit (Applied Biosystems™-A28007) following manufacturers’ instructions and used for droplet digital PCR experiments.

### Droplet digital PCR

For hsa-miR-221-3p detection in EVs and in cells, 1:200 cDNA was mixed with 1X hsa-miR-221-3p Advanced miRNA Assay (477981_mir, ThermoFisher-A25576) and ddPCR Supermix for Probes (Bio-Rad - 1863026) to a final volume of 23 μl. Y3 RNA was used as a reference. The same diluted cDNA was mixed with 50 nM Y3 primers (mouse: mY3 Fw: 5’-GGTTGGTCCGAGAGTAGTGG-3’, mY3 Rv: 5’-AAAGGCTGGTCAAGTGAAGC-3’; Human: hY3 Fw: GGCTGGTCCGAGTGCAGTG, hY3 Rv: GAAGCAGTGGGAGTGGAGAA) and QX200 ddPCR EvaGreen Supermix (Bio-Rad-1864033) to a final volume of 23 μl. Droplet formation and PCR conditions were performed following the manufacturers’ instructions using ddPCR™ 96-well plate (Bio-Rad). QX200 Droplet Reader Bio-Rad was used to read the plates. Analysis and target quantification were performed using QuantaSoft^TM^ Analysis Pro Software.

### Immunoblotting

Cells were lysed in lysis buffer (50C:mM Tris-HCl pH7.4, 150C:mM NaCl, 1 mM EDTA, 0.25% NP-40, 0.1% Triton X-100, 0.1% SDS, 1X protease inhibitor (ThermoFisher-78429). Cell lysates were loaded on 10% acrylamide gels and transferred to a PVDF membrane. Membranes were incubated with 1:1000 primary antibody and 1:10000 peroxidase-conjugated secondary antibodies in 3% milk in TBST. The signal was measured with Amersham ECL HRP-Conjugated Antibodies (Cytiva) using BioRad Chemidoc XRS+.

The following primary antibodies were used: anti-hnRNPA2B1 (PA5-34939, Invitrogen), anti-Calnexin (Abcam), anti-MycTag (Proteintech). The following secondary antibodies were used: goat Anti –Rabbit (Jackson ImmunoResearch Laboratories, Inc.), goat Anti Mouse (Jackson ImmunoResearch Laboratories, Inc.). Band areas and pixel intensities were quantified using ImageJ software.

### Absorbance scan and fluorescence indicator displacement (FID)

Five μM RNA EXO motif were mixed with 25 μM compounds in AlphaBuffer (25 mM HEPES pH 7.4, 100 mM NaCl, 0.01% BSA) to a final volume of 20 μL. Absorbance spectrum (200-530 nm) was measured using Tecan Spark® microplate reader with a NanoQuant plate at room temperature. In FID, a 2X final concentration of Midori green advanced (Resnova) was added to the samples and the melting curve measured using Bio-Rad CFX96™ System from 30 °C to 90 °C.

### Pull down (PD) assay

1-3*10^6^ NSC34 cells were lysed in buffer R-Lysis (25 mM HEPES pH 7.5, 100 mM NaCl, 1X protease inhibitor (ThermoFIsher, 78429)) and subjected to a water-bath sonication (35-40 amplitude, 6–7 cycles of 7 seconds on and 45 seconds off). After a centrifugation at 14000 rpm for 20 min at 4°C, the supernatant was collected and 200 µg were incubated with 5 μl of buffer equilibrated Dynabeads^TM^ M-280 Streptavidin (Invitrogen, 11205D) for 15 min at 4 °C in rotation to perform a pre-clearing step. After the magnetic separation, the pre-cleared lysate was incubated with 10 μl of 100 μM RNA biotinylated probes (or 400 ng of biotinylated EV-RNA) and incubated for 1h at 4 °C in rotation. Then, 5 μl/sample of beads were added and the sample were put again in rotation for 20 min at 4°C. After the magnetic separation the beads were washed with buffer R-Lysis (or 100 mM ammonium bicarbonate for proteomics) and bound proteins were eluted with 20 μl of 1X Laemmli sample buffer heating the samples at 95°C for 5 min, otherwise beads were resuspended in 40 μl with 100 mM ammonium bicarbonate for LC-MS analysis.

## LC-MS analysis

Proteins bound to the beads were subjected to on-bead trypsin digestion. Briefly, samples were reduced and alkylated with DTT 10 mM at 56°C for 45 min and iodoacetamide 20 mM at RT for 30 min in the dark, respectively. One microgram of trypsin (Thermofisher Scientific) was added to each sample and the beads were incubated at 37°C overnight with gentle shaking. Following digestion, beads were collected and the supernatant was transferred to a fresh Eppendorf tube. Beads were washed with 50 μl of 100 mM ammonium bicarbonate and the supernatants were pooled. Digested peptides were then acidified with 1% TFA to a pH 2.5, desalted on C18 stage-tips and resuspended in 20 μl of 0.1% formic acid buffer for LC-MS/MS analysis. Digested samples were separated using an Easy-nLC 1200 system (Thermo Scientific). A 28 cm reversed-phase column (inner diameter 75 µm packed in-house with ReproSil-Pur C18-AQ material: 3 µm particle size, Dr. Maisch, GmbH), heated at 40°C, was used for separating the peptides, with a two-component mobile phase system of 0.1% formic acid in water (buffer A) and 0.1% formic acid in 80% acetonitrile (buffer B). Peptides were eluted using a gradient of 5% to 25% over 57 min, followed by 25% to 40% over 13 min and 40% to 98% over 10 min, and kept at 98% over 10 min, a flow rate of 400 nl/min. Samples were injected in an Orbitrap Fusion Tribrid mass spectrometer (Thermo Scientific, San Jose, CA, USA) and data acquired in data-dependent mode (2100 V). Temperature of the ion transfer tube was set at 275°C. Full scans were performed in the Orbitrap at 120.000 FWHM resolving power (at 200m/z), 50 ms maximum injection time, and an AGC target of 1x10e6. A mass range of 350-1100 m/z was surveyed for precursors, with first mass set at 140 m/z for fragments. Each full scan was followed by a set of MS/MS scans (HCD, collision energy of 30%) over 3 sec cycle time at 150ms maximum injection time (ion trap) and AGC target of 5x10e3. A dynamic exclusion filter was set every 30 sec. Data were acquired using the Thermo software Xcalibur (version 4.3) and Tune (version 3.3). QCloud was used to control instrument longitudinal performance^78^.

Peptides searches were performed in Proteome Discoverer 2.2 software (Thermo Scientific) against the Mus musculus FASTA file (uniprot, downloaded April 2021) and a database containing major common contaminants. Proteins were identified using the MASCOT search engine, with a mass tolerance of 10 ppm for precursors and 0.6 Da for products. Trypsin was chosen as the enzyme with 5 missed cleavages. Static modification of carbamidomethyl (C) and variable modification of oxidation (M) and acetyl (protein N-term) were incorporated in the search. False discovery rate was filtered for <0.01 at PSM, at peptide and protein level. Results were filtered to exclude potential contaminants and proteins with less than two peptides. Protein–protein network analyses were generated by STRING 11.5 tool (http://string-db.org) using medium confidence.

### Immunofluorescence staining

Cells were washed twice with PBS without Ca^2+^/Mg^2+^ (LifeTechnologies) and fixed with 4% PFA in PBS for 10 min at room temperature, afterwards washed three times with PBS. Fixed cells were permeabilized for 10 minutes in 0.2 % Triton X solution and then incubated for 1 hour at RT in blocking solution (1% BSA, 5% donkey serum, 0.3M glycine and 0.02% Triton X in PBS). Primary antibodies were diluted in blocking solution and cells were incubated with primary antibody solution overnight at 4°C. The following primary antibodies were used: rabbit anti-Islet (1:500, Abcam), goat anti-ChAt (1:500, Millipore), mouse anti-MAP2 (1:500, BD Biosciences). Nuclei were counter stained using Hoechst (LifeTechnologies).

### Statistical analysis

MS downstream analysis were performed using the ProTN proteomics pipeline (www.github.com/TebaldiLab/ProTN and www.rdds.it/ProTN). Briefly, peptide intensities were log2 transformed, normalised (median normalisation) and summarised into proteins (median sweeping) with functions in the DEqMS Bioconductor package^79^. Imputation of the missing intensities was executed by PhosR package^80^. Differential analysis was performed with the DEqMS package, proteins with *p* value<0.05 were considered significant. Functional enrichment analysis of differentially expressed proteins was performed with EnrichR^81^. Enriched terms with *p* value<0.05 and overlap size>4 were considered significant. Additional statistical analyses were performed using *t*-test in the GraphPad Prism^®^ software, version 9.0. Comparisons were considered statistically significant if *p* values were <0.05 (*), <0.01 (**), or <0.001 (***).

