## Supplementary figures for "Small molecule inhibitors of hnRNPA2B1-RNA interactions reveal a predictable sorting of RNA subsets into extracellular vesicles"

**Supplementary Figures and Figure legends**

### Figure S1

**A**

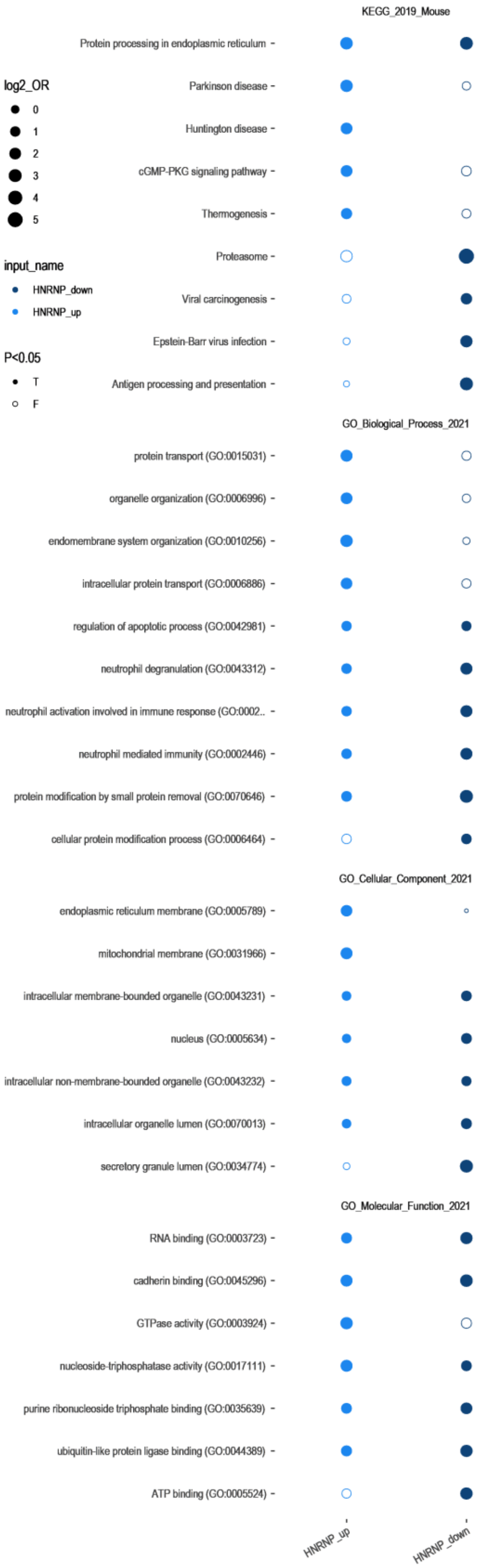


**B**

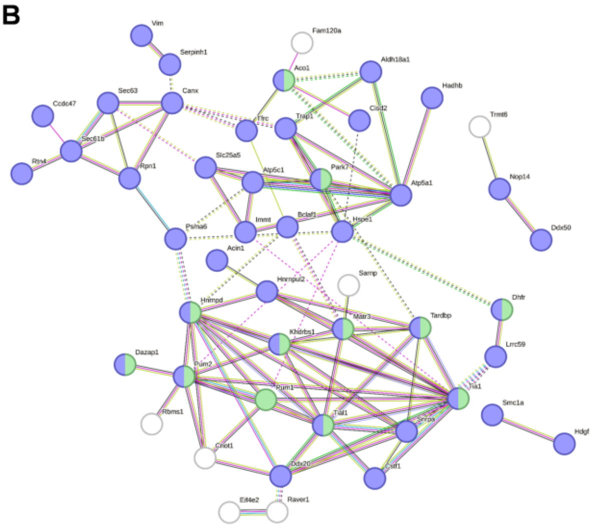

| compartment | description | count in network | strength | false discovery rate |
|---|---|---|---|---|
| Subcellular localization (COMPARTMENTS) | | | | |
| GOCC:0043231 | Intracellular membrane-bounded organelle | 49 of 6320 | 0.46 | 1.46e-14 |
| GOCC:0043229 | Intracellular organelle | 52 of 8073 | 0.38 | 3.68e-13 |
| GOCC:0005622 | Intracellular | 55 of 9943 | 0.32 | 2.44e-12 |
| GOCC:0070013 | Intracellular organelle lumen | 22 of 1992 | 0.62 | 1.34e-06 |
| GOCC:0036464 | Cytoplasmic ribonucleoprotein granule | 8 of 172 | 1.24 | 6.19e-06 |

| GO-term | description | count in network | strength | false discovery rate |
|---|---|---|---|---|
| Molecular Function (Gene Ontology) | | | | |
| GO:0003723 | RNA binding | 29 of 1034 | 1.02 | 1.98e-19 |
| GO:0003676 | Nucleic acid binding | 34 of 2889 | 0.64 | 1.85e-12 |
| GO:0003729 | mRNA binding | 13 of 270 | 1.26 | 4.85e-10 |
| GO:1901363 | Heterocyclic compound binding | 38 of 4771 | 0.47 | 1.76e-09 |
| GO:0097159 | Organic cyclic compound binding | 38 of 4858 | 0.47 | 2.52e-09 |
| GO:0003730 | mRNA 3-UTR binding | 7 of 79 | 1.52 | 1.64e-06 |
| GO:0005488 | Binding | 51 of 11199 | 0.23 | 4.30e-06 |
| GO:0035925 | mRNA 3-UTR AU-rich region binding | 3 of 27 | 1.62 | 0.0281 |
| GO:0035198 | miRNA binding | 3 of 30 | 1.57 | 0.0281 |
| GO:0051082 | Unfolded protein binding | 4 of 82 | 1.26 | 0.0281 |

**C**

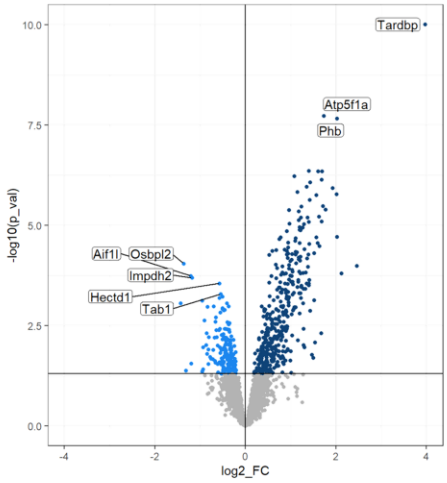


**D**

| Detected hnRNPs | Enriched RNA binding motifs | Common motifs |
|---|---|---|
| Hnrnpd | UGCAUUUAU<br>GUACUC<br>UUAUUUUAGUUAG | GAAGGG |
| Hnrnpc | AUAUUUUUAU | |
| Hnrnpa2b1 | UAGUGCCC<br>UGCAGAUGGUUAGUU<br>GGAAUUAAG | |
| Hnrnph2 | UGGGGA<br>GGGAAGAGC | |
| Hnrnpl | GGAAAA<br>ACCUAG | |
| Hnrnpa1 | AGAAAUGGA<br>UAGGGUUGU<br>CUGACC | |
| Hnrnph1 | GAAGGGGA<br>AGAGCA | |
| Hnrnpf | AAGGGGAGGGG<br>AGAGCAA | |
| Hnrnpk | UGUCAACCAG<br>AAAGAGAAUAAAGA | |

**Figure S1**. **Ontologies and RNA consensus motifs of hnRNPs identified by reverse proteomics**. **(A)** Gene Ontology (GO) analysis of proteins enriched (up) or displaced (down) in the ratio between hnRNPA2B1 OE and Mock conditions. The dimension of the point is given by the Odds Ratio in log2 scale. The filling is based on the significance of the adj. *p* value against the selected threshold. **(B)** Interaction network inside the 336 enriched protein targets obtained from STRING database (https://string-db.org). **(C)** Volcano Plot representing the differentially expressed proteins as described in Figure 1 but using lysates from TDP43-expressing cells. Raw data are available in Table S3. **(D)** Experimentally validated RNA sequence motifs reported as recognized by the identified hnRNPs Sequence motifs (http://rbpdb.ccbr.utoronto.ca) and analysed by MEME toolbox (https://meme-suite.org/meme).

**Figure S2**

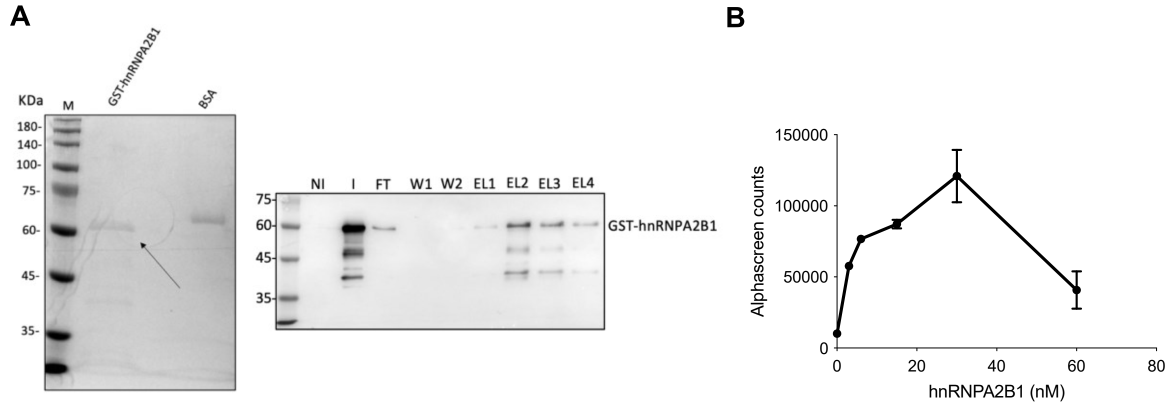


**Figure S2. GST-hnRNPA2B1 protein purification and AlphaScreen titration**. **(A)** Left: Coomassie of GST-hnRNPA2B1 purification. GST-hnRNPA2B1 concentration was calculated using 1 µg of gel-loaded BSA as reference. Right: Western blot using an anti-hnRNPA2B1 antibody and confirming the purification of the protein. “NI” non-induced bacteria; “I”, IPTG-induced bacteria, “W1, W2”, Wash 1 and 2; “EL”, elutions. **(B)** Graph showing the hooking point of the protein, reached at 30 nM. Hill coefficient was equal to 0.3 as calculated by nonlinear regression using specific binding with Hill slope fitting model of GraphPad Prism 9.

**Figure S3**

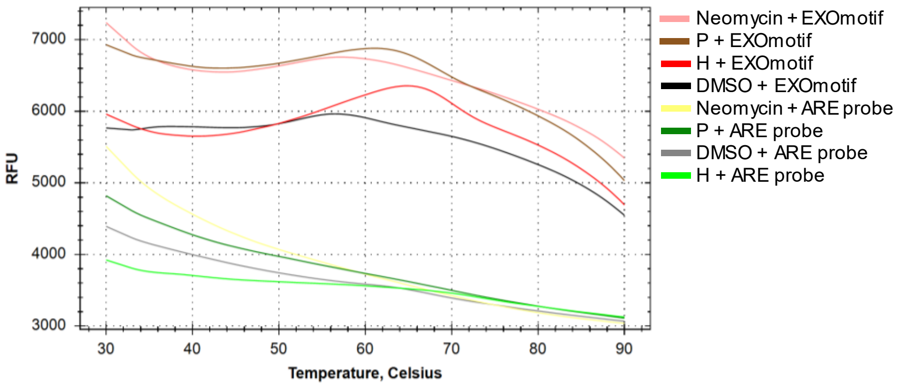


**Figure S3. Fluorescence intensity displacement (FID).** Purine-rich and ARE RNA probes were used for melting curve profiling using Midori green fluorescence in the presence of P and H compounds.

### Figure S4

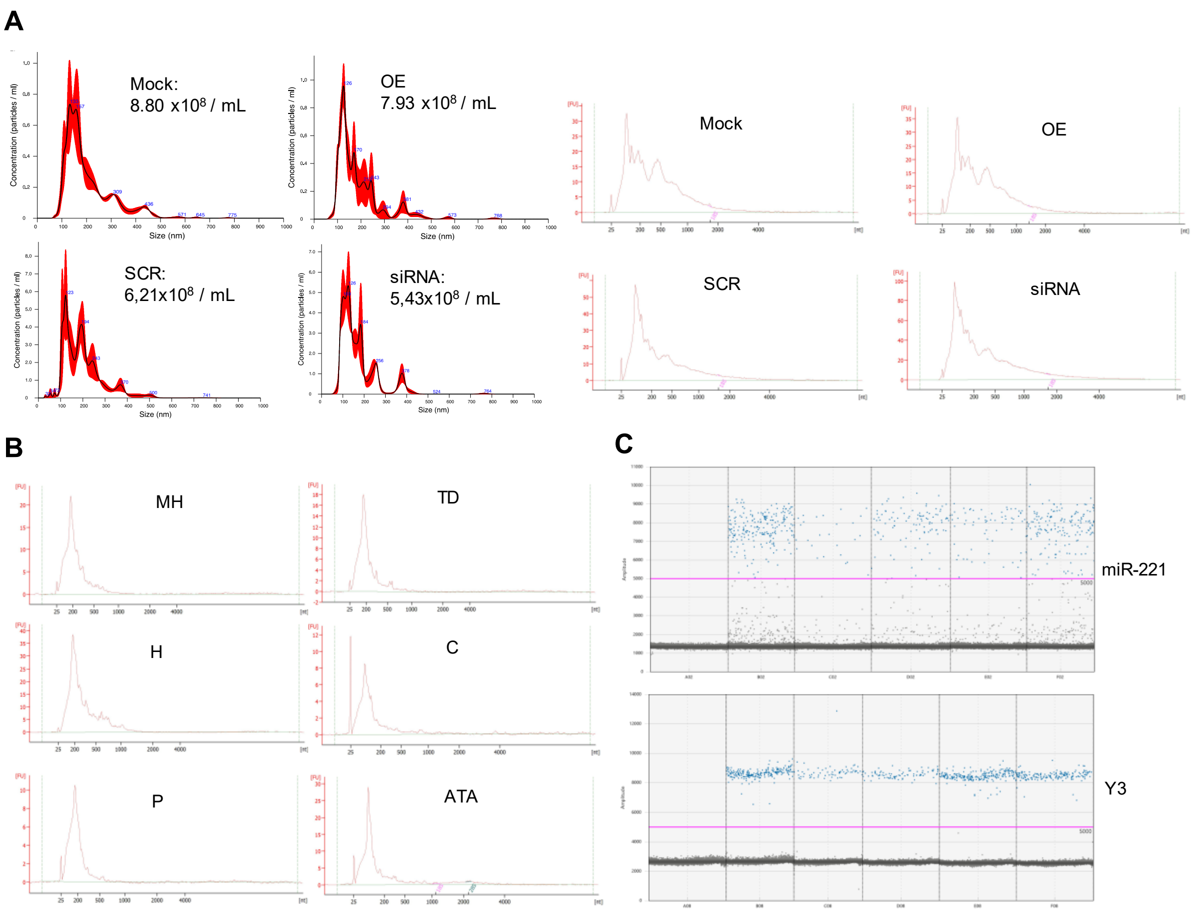


**Figure S4. EV-RNA analysis upon hnRNPA2B1 ectopic expression or silencing and compound treatment**. **(A)** Left: Representative NTA profiles of EVs retrieved after hnRNPA2B1 overexpression or silencing. Right: Representative Bioanalyzer electropherogram of EV-RNA retrieved after hnRNPA2B1 overexpression or silencing. **(B)** Representative Bioanalyzer electropherogram of EV-RNA retrieved after compounds treatment. **(C)** Representative ddPCR outputs relative to miR-221-3p and Y3 RNAs.

**Figure S5**

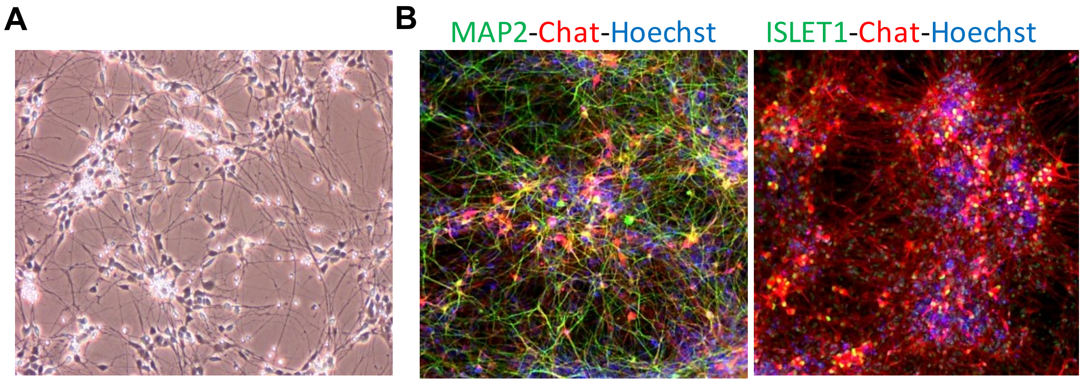


**Figure S5. Characterization of iPSC-derived motor neurons. (A)** Representative brightfield image of 10 days-differentiated and 2 weeks-matured motor neurons. **(B)** **I**mmunofluorescence stainings of iPSC-derived motor neurons after final differentiation. Left: pan neuronal marker microtubule associated protein tau (MAP2), mature lower motor neuron marker choline acetyltransferase (ChAt), total cell nuclei marker Hoechst. Right: Early motor neuron marker transcription factor ISELT1, mature lower motor neuron marker choline acetyltransferase (ChAt), total cell nuclei marker Hoechst. Magnification 20x.
